# Ecological factors rather than barriers to dispersal shape genetic structure of algal symbionts in horizontally-transmitting corals

**DOI:** 10.1101/037994

**Authors:** SW Davies, FC Wham, MR Kanke, MV Matz

## Abstract

Many reef-building corals acquire their algal symbionts (*Symbiodinium* sp.) from the local environment upon recruitment. This horizontal transmission strategy where hosts pair with locally available symbionts could serve to increase coral fitness across diverse environments, as long as hosts maintain high promiscuity and symbionts adapt locally. Here, we tested this hypothesis in two coral species by comparing host and symbiont genetic structures across different spatial scales in Micronesia. Each host species associated with two genetically distinct *Symbiodinium* lineages, confirming high promiscuity in broadly dispersing hosts. However, contrary to our initial expectation, symbiont genetic structure was independent of physical barriers to dispersal between islands, unlike genetic structure of their hosts that was nearly perfectly explained by ocean currents. Instead, *Symbiodinium* consistently demonstrated genetic divergence among local reefs and between the two host species at each island, although not necessarily between distant islands. These observations indicate that *Symbiodinium* lineages disperse much more broadly than previously thought and continuously adapt to specific hosts and reef environments across their range, following the classical Baas Becking’s hypothesis: “Everything is everywhere, but the environment selects”. Overall, our findings confirm that horizontal transmission could be a mechanism for broadly dispersing coral species to enhance their local fitness by associating with locally adapted symbionts. Dramatic differences in factors driving the genetic structures of horizontally-transmitting corals and their *Symbiodinium* imply that viewing their combined genomes as a single entity (‘hologenome’) would not be useful in the context of their evolution and adaptation.

## Introduction

Many well-known symbioses involve the passing of symbionts from parents to offspring (vertical transmission), fully aligning the evolutionary trajectories of symbiotic partners and typically lead to their deep integration at biochemical and genomic levels (i.e. Buchnera in aphids (1, 2)). The result of such symbiosis is essentially a novel composite organism. In other types of symbioses, the association between partners must be newly established in every generation (horizontal transmission), which allows for the maintenance of each partner’s species identity. In theory, this kind of relationship should generate novel ecological opportunities for both symbiotic partners through their mixing and matching across environments. For example, association with ecologically specialized algal photobionts can lead to distinct ecological guilds of lichens (3) or allow a fungal partner to expand its geographic range across a broader climatic envelope (4). Similarly, in aphids, association with various horizontally transmitted bacterial symbionts allows these insects to colonize novel host plants across climatic zones (5). Considering these and other examples of ecological adaptation based on varying symbiotic associations, it has been argued that the joint genomic content of symbiotic systems should be studied as a single unit of evolution, the ‘hologenome’ (6, 7). However, the usefulness of this concept depends on whether the evolutionary trajectories of both symbiotic partners are sufficiently aligned to present a unified target of selection (8). Here, we explore this question in the symbiosis between a horizontally transmitting reef-building coral and dinoflagellate algae of the genus *Symbiodinium* (9).

Association with *Symbiodinium* is obligatory for many coral hosts that rely on algal photosynthesis for energy, while the algae benefit from protected and light-exposed residence as well as inorganic nutrients and CO2 concentration regulatory mechanisms provided by the host (10-13). Given the obligatory nature of this symbiosis for the host, it is somewhat surprising that in the majority of coral species (∽85%) *Symbiodinium* are not transmitted vertically, but rather must be acquired by the juvenile coral from its local environment (14, 15). One possible explanation is that dispersal ranges of aposymbiotic coral larvae typically extend over hundreds of kilometers (i.e. (16), while the environmental variation corals must deal with exists on much smaller spatial scales: reef environments with varying light, thermal and nutrient conditions can be separated by meter-scale distances (i.e. (17). Under such circumstances, coral hosts would benefit from the mixing and matching strategy, improving their fitness by associating with the locally available, and putatively ecologically specialized, algal strains (18-20). Establishing the relative roles of these symbiotic partners in adaptation to variable environments is essential for better prediction of coral reefs’ future under climate change (i.e. 21).

The working hypothesis for this study, the ‘global host, local symbiont’ hypothesis, posits that (i) coral host disperses widely and is able to establish symbiosis with diverse algal strains across locations; (ii) *Symbiodinium* algae are poor dispersers (i.e., 21), which results in strong divergence among locations aligning with physical barriers and facilitates their local environmental specialization; (iii) local *Symbiodinium* strains also diverge with respect to the host species as a result of selective pressure towards higher host specificity (22, 23). To validate all three components of our hypothesis, we examined multi-locus genotypes (MLG) of clade C Symbiodinium in two species of *Acropora – A. hyacinthus* and *A. digitifera* – collected from the same reef locations across the Micronesian Pacific (Fig. 1A). Our previous work has shown that both host species exhibit extensive genetic connectivity and their genetic structure is nearly perfectly explained by the patterns of regional surface currents (16). By using two coral species that co-occur across the same locations as well as local reef environments we aimed to disentangle the roles of environmental specialization, host specialization, and physical barriers to dispersal in driving the fine-scale genetic structure of *Symbiodinium*.

**Figure 1:**
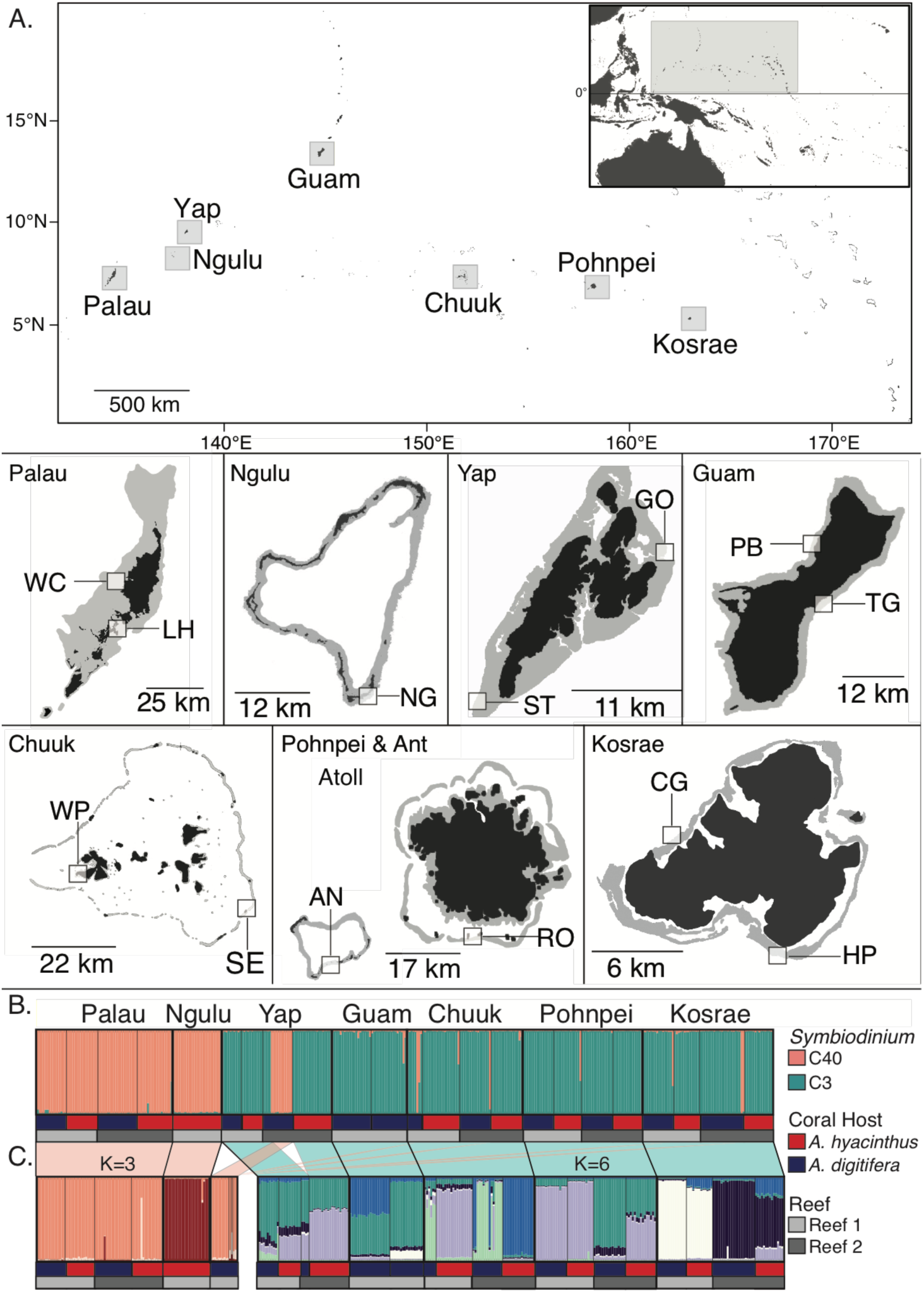
Geographic location of the Micronesian islands where *Acropora hyacinthus* and *A. digitifera* coral samples were collected (A). Top: Map of the Micronesian Pacific with an inset of the Pacific Ocean for reference. Islands where samples were collected and analyzed for *Symbiodnium* genetics are designated with grey boxes. Detailed information on each sampling site is located in Table S3. No *A. hyacinthus* were present in Guam and *A. digitifera* were not collected in Ngulu. Analyses of microsatellite loci data for *Symbiodinium* hosted by *Acropora hyacinthus* and *Acropora digitifera* using multilocus genotyping data. B. structure population assignment for *Symbiodinium* from two *Acropora* host species across greater Micronesia at an optimal population number (K=2), corresponding to C40 (pink) and C3 (turquoise) clade C subclades. Colors in the bottom panels correspond to host species and shades of grey correspond to different sites within each island. C. Within-subclade STRUCTURE analysis.

## Results

### Two *Symbiodinium* clade C lineages

To enable standard population genetic analysis, data analysis was restricted to corals hosting a single diploid *Symbiodinium* clone, which encompassed the majority (69% *A. digitifera* and 64% *A. hyacinthus*) of our samples. Across the two coral species in Micronesia (Fig. 1A), two distinct *Symbiodinium* lineages were observed, which were most clearly discriminated by two of the six microsatellite loci (Fig. S1). Sequencing of the psbA_ncr_ gene for a subset samples classified these lineages as C40 and C3 (39, (24) (Fig. S2). Corals of both species from Palau and Ngulu hosted exclusively C40 (Fig. 1B, pink bars). This lineage was also prevalent in *A. digitifera* at one reef site on Yap and was rarely found in *A. digitifera* across the rest of Micronesia. All other coral hosts of either species associated with C3 (Fig. 1B, turquoise bars). Both *Symbiodinium* lineages possessed high genetic diversity, with a total of 70 unique alleles across three islands in C40 (Table S1A) and 130 unique alleles across five islands in C3 (Table S1B). Mean numbers of alleles per island for each locus for C40 ranged from 3.00-4.67 with the highest numbers of private alleles in Palau (N=7) (Table S1A). Mean allele numbers per island for each locus for C3 ranged from 3.83-5.17 and numbers of private alleles ranged from 1-4 (Table S1B).

### *Symbiodinium* genetic structure

All pairwise between-island *F*_ST_’s for C40 were significant (Table 1A). C3 had one non-significant *F*_ST_ (Yap-Pohnpei) while all others were significant and ranged from 0.058 (Guam-Chuuk) to 0.078 (Yap-Kosrae) (Table 1B). Notably, *Symbiodinium* C3 differentiation did not show the isolation-by-distance (IBD) pattern observed in both coral hosts (Fig. 2). IBD could not be computed for C40 due to too few between-island comparisons. C3 *F*_ST_ values generally exceeded *F*_ST_ values for the host (Fig. 2A), however, this result did not imply stronger symbiont genetic differentiation: higher *F*_ST_ in the symbionts was a consequence of higher mean heterozygosity in *Symbiodinium* markers (e.g. (25). Indeed, an alternative measure of genetic differentiation controlling for mean heterozygosity - Jost’s D (26) - demonstrated that symbiont genetic divergence was in fact intermediate between the two hosts (Fig. 2B).

**Table 1.** Summary of pairwise *F*_ST_ values between all islands for *Symbiodinium* C40 (A) and C3 (B). Permutations were run 9999 times. All significant comparisons are shaded in grey.

|  | PAL | NGU | YAP |
| --- | --- | --- | --- |
| PAL | 0.000 | *** | 0.004 |
| NGU | 0.279 | 0.000 | *** |
| YAP | 0.056 | 0.326 | 0.000 |

B. *Symbiodinium* C3
|  | YAP | GUA | CHU | POH | KOS |
| --- | --- | --- | --- | --- | --- |
| YAP | 0.000 | *** | *** | 0.059 | *** |
| GUA | 0.062 | 0.000 | *** | *** | *** |
| CHU | 0.066 | 0.058 | 0.000 | *** | *** |
| POH | 0.009 | 0.071 | 0.063 | 0.000 | *** |
| KOS | 0.078 | 0.077 | 0.076 | 0.067 | 0.000 |

**Figure 2:**
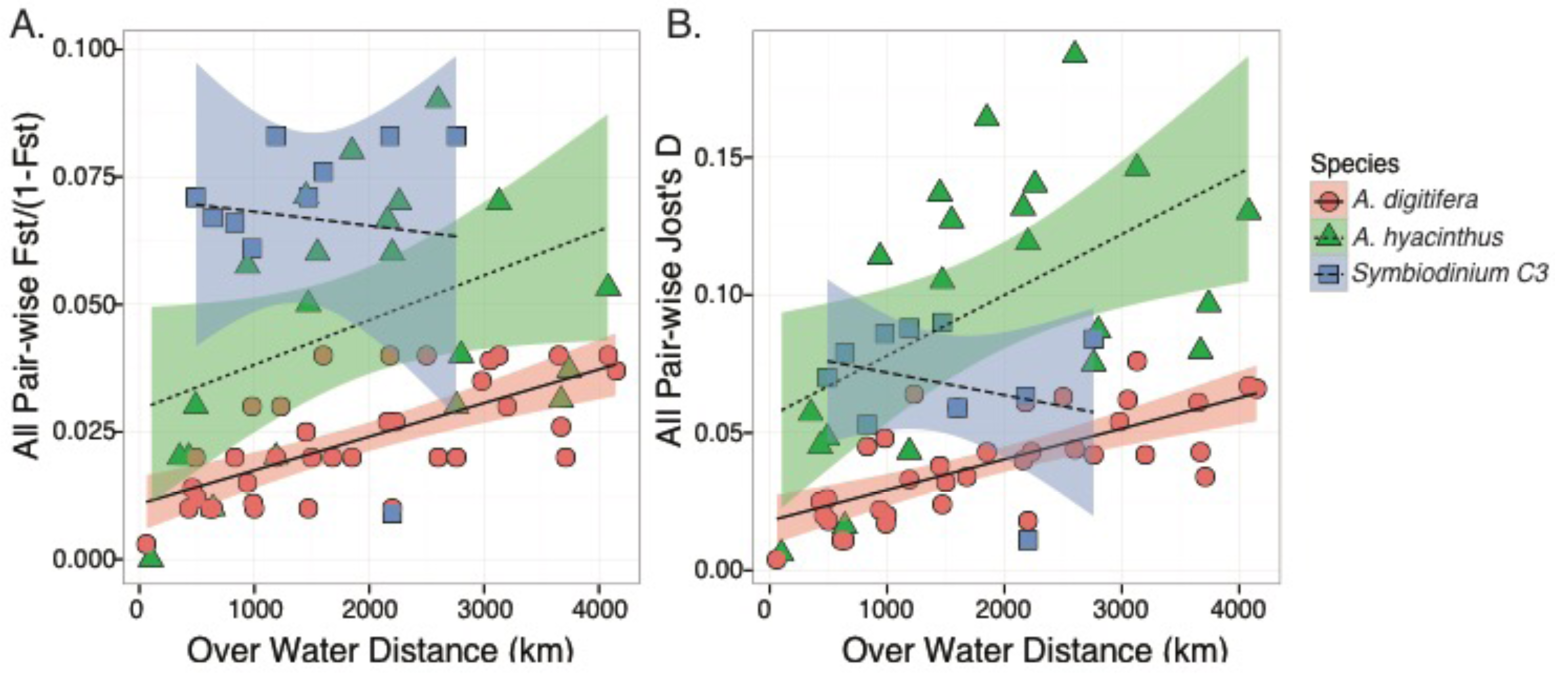
Comparison of two host species and *Symbiodinium* C3 differentiation. A. Pairwise genetic differentiation [(*F*_ST_ /(1-*F*_ST_)] of two species of *Acropora* coral and *Symbiodinium* C3 across linear distances (km) demonstrating significant isolation by distance for the two host species but no correlation for the symbiont. B. Pairwise Jost’s D for the same two host species and *Symbiodinium* C3 across linear distances (km) demonstrating no isolation by distance and no difference in overall divergence between the host and symbiont.

### Host specificity and environmental specialization

Discriminant analysis of principal components (DAPC) strongly differentiated between host species for both *Symbiodinium* C40 and C3 (Table S2, Fig. S3A, B), suggesting that host specificity drives symbiont diversity. In addition, DAPC demonstrated consistent differences among islands for each *Symbiodinium* lineage irrespective of host species: strong per-island assignment proportions were observed for C40 (Fig. S3C, 82-100%) and C3 (Fig. S3D, 51-98%), consistent with both STRUCTURE (Fig. 1B) and *F*_ST_ results (Table 1). Moreover, nearly all pairwise *F*_ST_ values between different reef sites within islands were significant for both lineages (Table 2), suggesting environmental partitioning of symbionts. In accord with these results, of the two top eigenvalues in DAPC of individual within islands, one explained divergence by host species (host specificity) and the other explained differences between reef sites (environmental specialization) (Fig. 3).

**Table 2.** Summary of pairwise *F*_ST_ values for *Symbiodinium* C40 (A) and C3 (B) between all sites pooled across host species. Permutations were run 9999 times. All significant comparisons are shaded in grey.

A. *Symbiodinium* C40
|  | PAL1 | PAL2 | NGU | YAP2 |
| --- | --- | --- | --- | --- |
| PAL1 | 0.000 | 0.009 | *** | 0.007 |
| PAL2 | 0.031 | 0.000 | *** | 0.007 |
| NGU | 0.283 | 0.303 | 0.000 | *** |
| YAP2 | 0.057 | 0.074 | 0.326 | 0.000 |

B. *Symbiodinium* C3
|  | YAP1 | YAP2 | GUA1 | GUA2 | CHU1 | CHU2 | POH1 | POH2 | KOS1 | KOS2 |
| --- | --- | --- | --- | --- | --- | --- | --- | --- | --- | --- |
| YAP1 | 0.000 | 0.139 | 0.002 | 0.009 | *** | *** | 0.002 | 0.055 | *** | *** |
| YAP2 | 0.011 | 0.000 | *** | 0.003 | *** | *** | 0.027 | 0.064 | *** | *** |
| GUA1 | 0.059 | 0.107 | 0.000 | 0.045 | *** | *** | *** | *** | *** | *** |
| GUA2 | 0.045 | 0.062 | 0.025 | 0.000 | 0.002 | *** | *** | 0.008 | *** | *** |
| CHU1 | 0.071 | 0.073 | 0.104 | 0.065 | 0.000 | *** | *** | *** | *** | *** |
| CHU2 | 0.079 | 0.112 | 0.070 | 0.086 | 0.068 | 0.000 | *** | *** | *** | *** |
| POH1 | 0.053 | 0.028 | 0.126 | 0.111 | 0.085 | 0.126 | 0.000 | *** | *** | *** |
| POH2 | 0.018 | 0.018 | 0.093 | 0.039 | 0.067 | 0.092 | 0.063 | 0.000 | *** | *** |
| KOS1 | 0.154 | 0.146 | 0.170 | 0.110 | 0.163 | 0.176 | 0.168 | 0.119 | 0.000 | *** |
| KOS2 | 0.092 | 0.121 | 0.116 | 0.131 | 0.129 | 0.090 | 0.136 | 0.100 | 0.183 | 0.000 |

**Figure 3:**
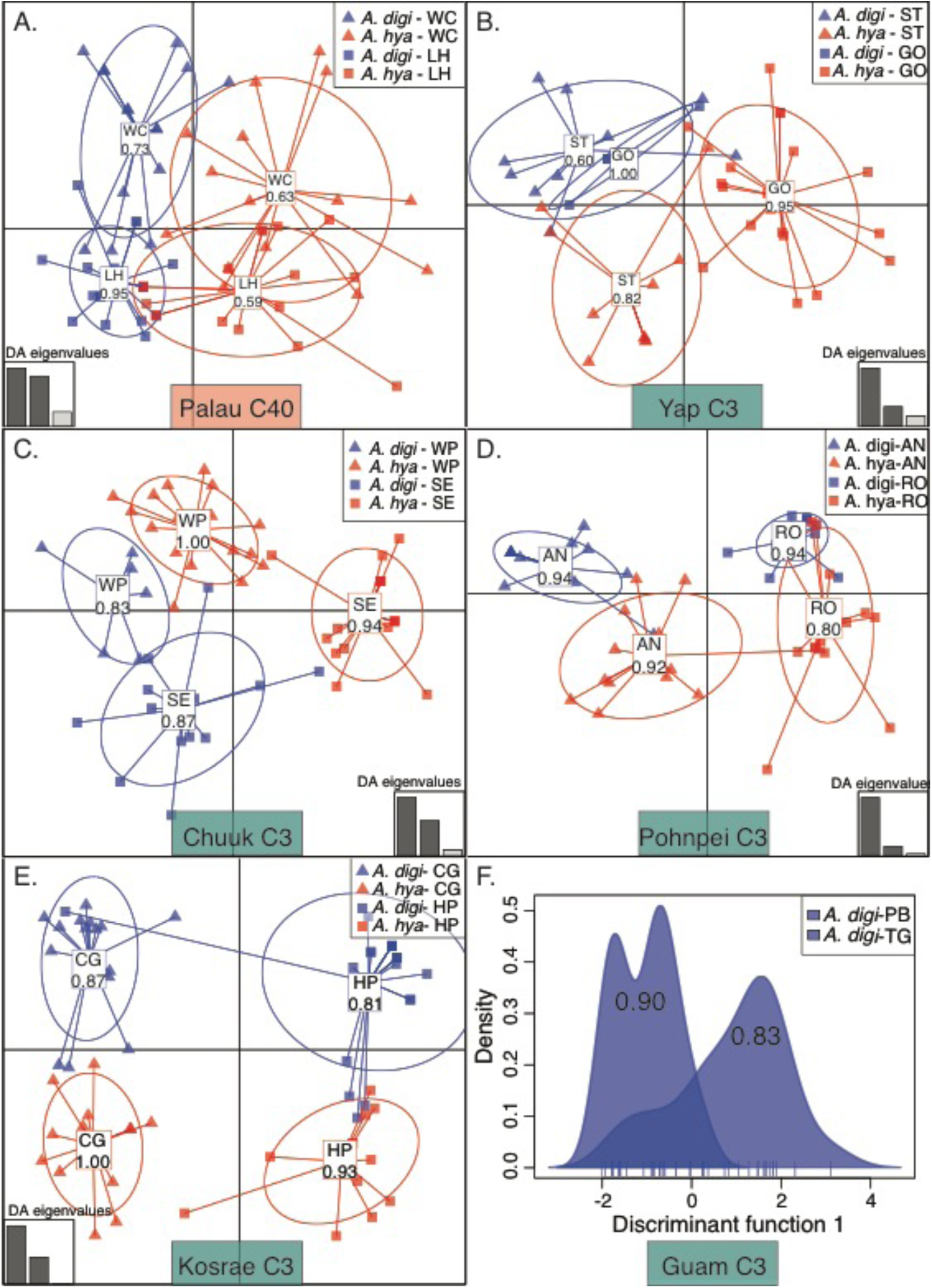
Discriminant analysis of principal components (DAPC) of MLG data for *Symbiodinium* C40 and C3 hosted by *Acropora hyacinthus* and *Acropora digitifera* at twelve sites across six islands (Ngulu not included) in Micronesia. DAPC analysis on two discriminant functions demonstrating host species assignments and site assignments. DAPC scatter plots for individual samples from within A. Palau for *Symbiodinium* C40. B. Yap for *Symbiodinium* C3, C. Chuuk for *Symbiodinium* C3, D. Pohnpei for *Symbiodinium* C3, E. Kosrae for *Symbiodinium* C3, and F. Guam for *Symbiodinium* C3 for *A. digitifera* hosts only (1 DF axis). Proportions of assignments are indicated in the clusters. Information on the DAPC models can be found in Table S2.

## Discussion

### Promiscuity of the coral host

Clade C *Symbiodinium* are considered to be the most derived lineage within the genus *Symbiodinium* and exhibit significantly higher within-clade diversity when compared to other, more basal, clades (9, 27, 28). Across the Micronesian Pacific (Fig. 1A), both coral hosts associated exclusively with clade C *Symbiodinium*, which was nevertheless represented by two distinct lineages– C40 and C3 (Fig. S2). This observation confirms the first prediction of the ‘global host, local symbiont’ hypothesis: coral hosts show considerable flexibility in their symbiotic association across their range and within their habitat.

### *Lack of dispersal limitation in* <u>Symbiodinium</u> *across Micronesia*

Initially, we expected to find strong isolation by distance in *Symbiodinium*, since the prevailing view of the *Symbiodinium* life cycle involves symbiotic existence in sedentary hosts alternating with a free-living form that largely exists in the benthos where dispersal by ocean currents must be limited (29-32). In contrast, we observed that genetic differentiation between islands of *Symbiodinium* C3 across Micronesia did not exceed *F*_ST_ = 0.078, which contrasts studies from other locations reporting symbiont *F*_ST_ as high as 0.54 in clade A (33) and Φ_ST_ as high as 0.468 in clade C (34). This lack of physical dispersal limitation is best demonstrated by comparing *Symbiodinium* C3 divergence to divergence of their coral hosts, which has been shown to nearly perfectly follow biophysical barriers (16). If *Symbiodinium* dispersal were limited by the same physical barriers and to a greater extent, as our hypothesis posited, we would have observed steeper increases in divergence with distance relative to the host; instead, no increase of *F*_ST_ with distance was observed (Fig. 2A). Moreover, after correcting for mean heterozygosity, genetic divergence (Jost’s D) in *Symbiodinium* C3 was found to be intermediate between *A. hyacinthus* and *A. digitifera* (Fig. 2B), once again demonstrating that symbiont dispersal is less physically limited than their hosts. This finding contradicts the second prediction of our ‘global host, local symbiont’ hypothesis and is contrary to our expectations based on earlier *F*_ST_-based reports of stronger *Symbiodinium* differentiation compared to hosts (20-22, 34).

### Host specificity

The majority of reef-building coral species associate with a specific strain (termed “clade” or “subclade”) of *Symbiodinium* broadly defined based on ribosomal and/or chloroplast markers (22, 35-37). Previous *Symbiodinium* multilocus genotyping studies revealed that each of these strains harbors greater diversity, both in terms of genetic and functional diversity (20, 34, 38). Our data indicate that the local association of hosts and symbionts of the same genotypic cluster is due to pervasive evolution of host specificity in *Symbiodinium* (Fig. S3 & 3). As our study includes two coral species, we also observe that this specificity is not perfect: at every location there were symbionts that would have been assigned to another coral host based on their multilocus genotype (Fig. 3). This suggests that host specialization in Symbodinium arises in the face of considerable exchange between symbiont communities hosted by different coral species at the same location.

### Environmental partitioning

The most ecologically relevant postulate of the ‘global host, local symbiont’ hypothesis is that locally available symbionts are also locally adapted, giving rise to a locally adapted holobiont. Recurrent genetic divergence of symbionts between reef sites within the same island (Fig. 3) despite the lack of physical dispersal limitation across much larger distances (Fig. 2) suggests that symbiont genetic divergence is likely due to poor survival of immigrants rather than to physical barriers to migration. Given that hosts are available at every site, this leaves other ecological parameters of local reef environment as the most likely barrier-forming force, preventing survival of immigrants adapted to a different environment– a situation termed “phenotype-environment mismatch” (39). This in turn implies environmental specialization (i.e., local adaptation) in the symbionts, which supports our hypothesis. Interestingly, just like host specificity, this environmental specialization is imperfect: there are several *Symbiodinium* genotypes that appear to be successful migrants between reef sites (particularly clearly visible for Kosrae, Fig. 3E), indicating that environmental partitioning also arises in the face of considerable gene flow.

## Conclusions

Both coral host species associated with two divergent *Symbiodinium* lineages, indicating that coral hosts are promiscuous in their symbiont associations across the seascape. In contrast, within each *Symbiodinium* lineage, strong associations with particular host species were observed, suggesting that host-specificity is an important driver of *Symbiodinium* diversification. Unexpectedly, *Symbiodinium* genetic divergence did not correlate with physical distance between islands, but was driven by the combined effects of local (within-island) environment and host identity. This indicates that *Symbiodinium* assemblages across Micronesia are comprised by broadly dispersing lineages dynamically adapting to specific coral hosts and reef environments across their range. Assuming that host species can be regarded as part of *Symbiodinium*’s environment, this pattern clearly follows the classical hypothesis of Baas Becking (originally proposed for soil microbes): “Everything is everywhere, but the environment selects” (40). Overall, our observations support the view that horizontal transmission might be a strategy for broadly dispersing coral hosts to improve their local fitness by associating with locally available, environmentally specialized, *Symbiodinium*. Finally, dramatic differences in factors that structure genetic diversity in the coral host and *Symbiodinium* indicate that viewing their combined genomes as a single ‘hologenome’ would not be useful in the context of their evolution and adaptation.

## Materials and Methods

### Sampling

This study comprised a subset of samples previously analyzed for coral host genetics in Davies *et al.* (16) (Table S3, Fig. 1A). Twenty-five individuals of each coral host species (*Acropora hyacinthus* and *A. digitifera*) were examined at two reef sites within seven islands, with the exception of Ngulu (only collected *A. hyacinthus*) and Guam (no *A. hyacinthus* found), for a total of thirteen sites.

### Laboratory Procedures

Holobiont DNA was isolated following Davies *et al*. (41). Microsatellite primers consisted of six previously published clade C loci (42, 43) and one novel locus mined using Msatcommander (44) from nucleotide EST data for *Symbiodinium* sp. clade C3 (45) (Table S4). Loci were multiplexed in 20 μl polymerase chain reaction (PCR) mixtures containing 10 ng of template, 0.1 μM of each forward and reverse primers, 0.2 mM dNTP, 1 μl 10X *ExTaq* buffer, 0.025 U *ExTaq* Polymerase and 0.0125 U *Pfu* Polymerase. Amplifications began at 94°C for 5 min, followed by 35 cycles of 94°C for 40 s, annealing temperature for 120 s, and 72°C for 60 s and a 10 minute 72°C extension period. Molecular weights were analyzed using the ABI 3130XL capillary sequencer. Data were binned by repeat size and individuals failing to amplify at ≥3 loci were excluded from analyses.

### Data Analysis

*Symbiodinium* were scored as diploid based on available information on their ploidy (22, 43, 46) Since each host could potentially contain populations of genetically distinct *Symbiodinium*, multi-locus genotypes (MLG) were only considered if they contained two or fewer alleles across all loci, suggesting that all alleles originate from a single, clonally replicated, genome (46), which was the case for 190 of 277 *A. digitifera* hosts and 178 of 278 *A. hyacinthus* hosts. structure v2.3.3 (47) assigned similar MLGs to populations (10^6^ iterations, burn in = 300,000) across ten replicate runs for each K (1-10) using an admixture model with no location prior. Following (48), ΔK was calculated in structure Harvester (49) and clumpp (50) and distruct (51) produced all graphics. We observed high percent assignment to two clusters among sympatric individuals. Following (46), we investigated the relationship between these assignments and allele identities and found strong relationships between assignments and allele identity, particularly at Sgr_21 and Spl_33 (Fig. S1), and we concluded that two distinct *Symbiodinium* lineages were present. MLG data were then split into lineages and additional structure runs were completed.

Partitioned lineage data were investigated for genetic divergences among islands and sites using *F*_ST_ (AMOVA, 9999 permutations in genalex v6.5, (52). Pairwise differentiations (*F*_ST_ and Jost’s D) and pairwise island distances were correlated to test for isolation by distance (IBD) in *Symbiodinium* C3 (insufficient C40 data) and these trends were compared to results from the host species (53). Mean allelic diversities per island and numbers of private alleles per island were also calculated in genalex v6.5.

To resolve differences between islands, between host species and between sites and host species within each island, assignment of samples to genetic clusters using discriminant analysis of principal components (DAPC) was performed in R (54) using the ADEGENET package (55, 56). Data were converted into principle components (PCs) and then a-scores were used to determine trade-offs between power of discrimination and model over-fitting. Relationships were examined by DAPC and model information are listed in Table S2.

## Sequencing Analysis of *Symbiodinium* psbA^ncr^

To confirm individual assignments to different *Symbiodinium* lineages, the non-coding region of the circular plastid (psbA^ncr^) was amplified in a subset of samples using the methods described by (24). Amplified products were directly sequenced and aligned with ClustalW2 (http://www.ebi.ac.uk/Tools/msa/clustalw2/) to clade C psbA^ncr^ sequences from common Indo-Pacific *Symbiodinium* types (Table S5). The phylogeny was reconstructed using the default settings of Phylogeny.fr (57) and identities of *Symbiodinium* types were interpreted based on percent sequence identity with previously identified *Symbiodinium* types.

## Acknowledgements

Thanks to field assistants Carly Kenkel, Tim Kiett, Irina Yakushenok and David Stump. Nida Khawaja Rahman was integral in molecular work and James Derry assited with data management. We are grateful to Ulrich Mueller for numerous suggestions. We acknowledge all permit authorities. This study was supported by the Coral Reef Conservation program of the National Oceanic and Atmospheric Administration (administered through the Hawaiian Undersea Research Laboratory) and the National Science Foundation grant DEB-1054766 to M.V.M. DW was funded in part by Pennsylvania State University and grants from the National Science Foundation (OCE-0928764 and IOS-1258058).

## Supporting Information

**Figure S1:**
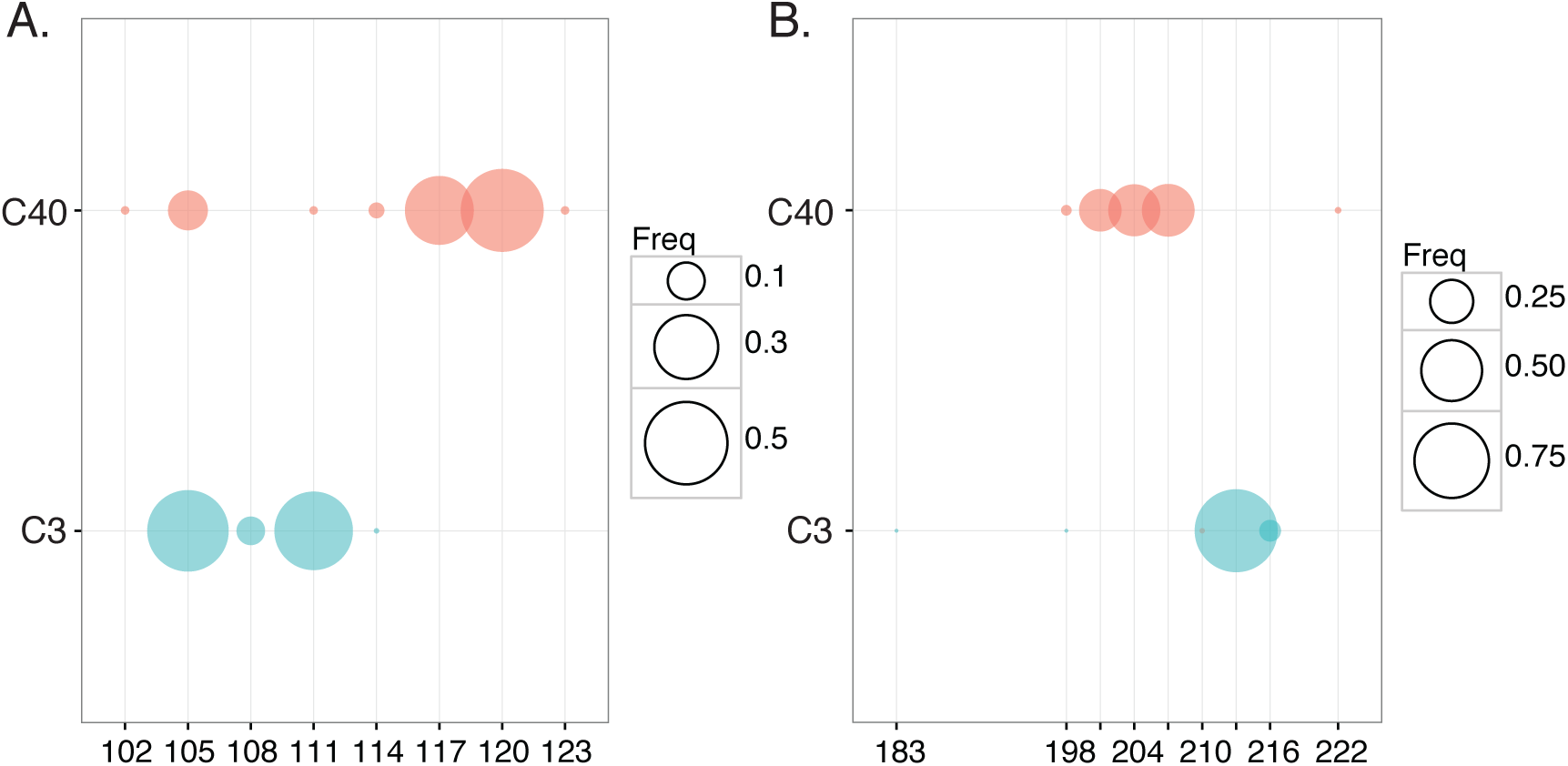
Bubble plot of the SSR allele frequencies of C3 (red) and C40 (blue) at the two loci that showed the largest discrimination power, Sgr_21 (A) and Spl_33 (B). The size of each bubble is proportional to the allele frequency within each *Symbiodinium* type.

**Figure S2.**
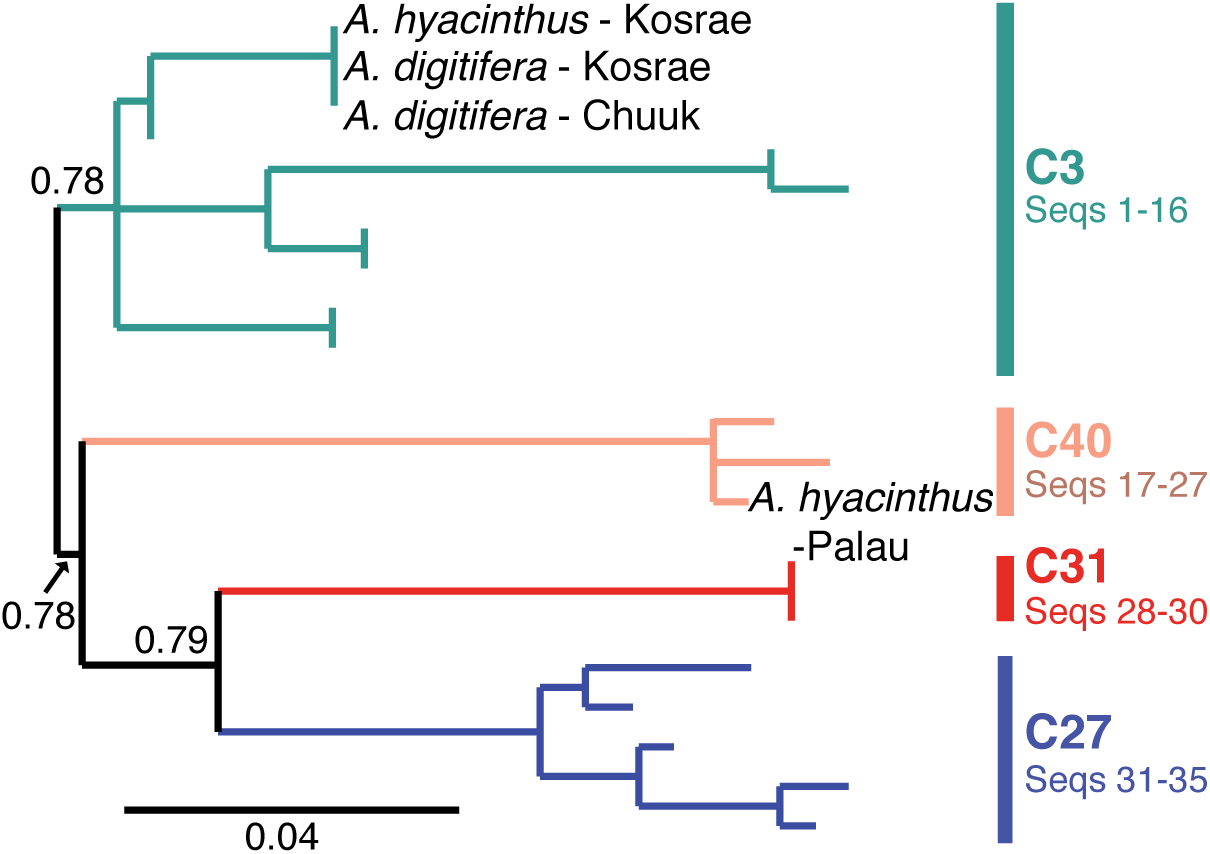
Phylogenetic analysis of psbA^ncr^ sequences from representative samples from this study (labeled branch tips) along with publically available psbA^ncr^ sequences from known *Symbiodinium* subclades identifies two lineages present (C3 and C40). Bootstrap support values are shown at the partitions that define known subclades. Scale bar: replacements per nucleotide site. Sequence accession numbers for reference sequences are given in Table S1.

**Figure S3:**
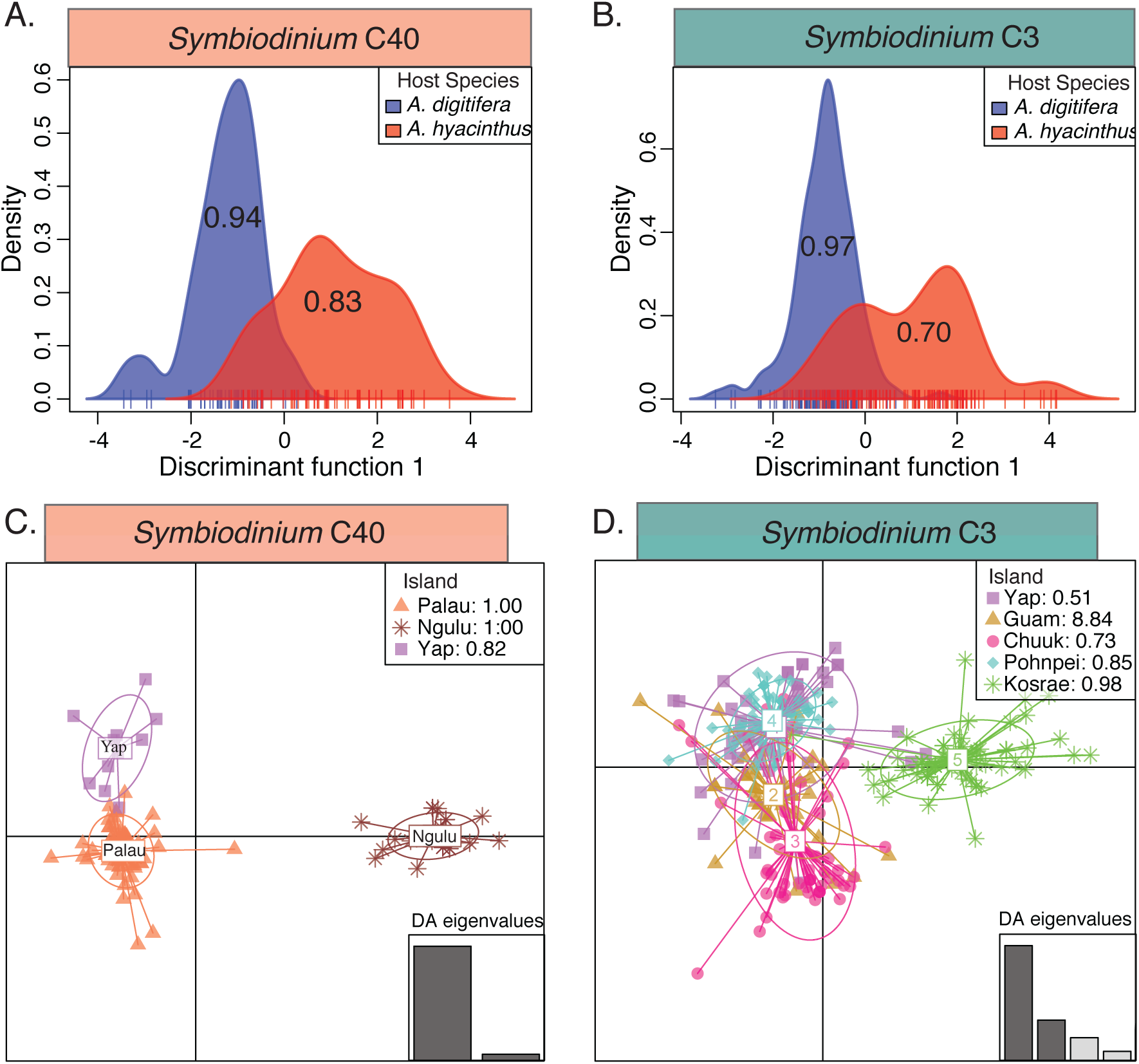
Discriminant analysis of principal components (DAPC) of MLG data for *Symbiodinium* C40 and C3 hosted by *Acropora hyacinthus* and *Acropora digitifera* at thirteen sites across seven islands in Micronesia. A. DAPC analysis on two discriminant functions demonstrating strong host species assignments across all islands for *Symbiodinium* C40 and B. *Symbiodinium* C3. Numbers overlaying the curves indicate successfully assigned fraction of samples. C. DAPC scatter plot for individual samples from *Symbiodinium* C40 represented by colored dots clustered by islands. D. DAPC scatter plot for individual samples from *Symbiodinium* C3 represented by colored dots clustered by islands. Proportions of assignments are indicated in the clusters or in the legends. Information on the DAPC models can be found in Table S2.

**Table S1:** Summary of allelic diversity measures for *Symbiodinium* C40 (A) and *Symbiodinium* C3 (B). Na: number of alleles, He: expected heterozygosity, Ho: observed heterozygosity, PA: number of private alleles

|  | Na | He | Ho | PA |
| --- | --- | --- | --- | --- |
| PAL | 4.67 | 0.52 | 0.64 | 7 |
| NGU | 4.00 | 0.42 | 0.55 | 6 |
| YAP | 3.00 | 0.43 | 0.51 | 2 |

B. *Symbiodinium* C3
|  | Na | He | Ho | PA |
| --- | --- | --- | --- | --- |
| YAP | 4.33 | 0.47 | 0.56 | 1 |
| GUA | 3.83 | 0.42 | 0.46 | 3 |
| CHU | 5.17 | 0.58 | 0.51 | 3 |
| POH | 4.33 | 0.47 | 0.56 | 1 |
| KOS | 4.00 | 0.48 | 0.51 | 4 |

**Table S2:** Discriminant analysis of principle component (DAPC) model information including the number of principle components (“PC”) and discriminant functions (“DF”) retained, the proportion of conserved variance by the clustering model (“var”), and the overall assignment proportions across the model (“assign”). A. DAPC information for *Symbiodinium* from different host species, B. islands, and C. sites and host species within each island.

| A. Host Species | PC | DF | var | assign |
| --- | --- | --- | --- | --- |
| *Symbiodinium* C40 | 7 | 1 | 0.814 | 0.879 |
| *Symbiodinium* C3 | 14 | 1 | 0.922 | 0.846 |
| B. Islands | PC | DF | var | assign |
| *Symbiodinium* C40 | 16 | 4 | 0.970 | 0.981 |
| *Symbiodinium* C3 | 19 | 4 | 0.966 | 0.796 |
| C. Within Island | PC | DF | var | assign |
| Palau C40 | 14 | 3 | 0.984 | 0.735 |
| Yap C3 | 7 | 3 | 0.844 | 0.778 |
| Guam C3 | 3 | 3 | 0.647 | 0.860 |
| Chuuk C3 | 12 | 3 | 0.933 | 0.927 |
| Pohnpei C3 | 8 | 3 | 0.906 | 0.900 |
| Kosrae C3 | 7 | 3 | 0.878 | 0.888 |

**Table S3:** Reef Site Collections. GPS coordinates, main island group, number of *A. digitifera* and *A. hyacinthus* hosts genotyped. The numbers in brackets are individuals hosting a single *Symbiodinium* MLG, which were included in all analyses presented here. Site letter corresponds to island insets in Figure 1A.

| Site | Island | GPS | *A. digitifera* | *A. hyacinthus* |
| --- | --- | --- | --- | --- |
| WC. West Channel Reef | Palau | 7°31'55.7 N, 134°29'42.8 E | 24 (20) | 25 (17) |
| LH. Lighthouse Reef | Palau | 7°16'62.4 N, 134°27'61.9 E | 23 (15) | 24 (16) |
| NG. Ngulu | Ngulu Atoll | 8°18'12.0 N, 137°29'18.7 E | 0<sup>1</sup> | 39 (25) |
| ST. South Tip Reef | Yap | 9°26'05.4 N, 138°02'10.4 E | 25 (10) | 25 (11) |
| GO. Goofnuw Channel | Yap | 9°34'26.4 N, 138°12'19.2 E | 24 (15)<sup>m</sup> | 25 (20) |
| PB. Pago Bay | Guam | 13°25'66.6 N, 144°47'94.3 E | 26 (20) | 0* |
| TG. Tanguisson | Guam | 13°32'61.1 N, 144°48'52.6 E | 21 (17) | 0* |
| WP. West Polle | Chuuk | 7°19'69.7 N, 151°33'21.1 E | 15 (6)<sup>m</sup> | 24 (18) |
| SE. South East Pass | Chuuk | 7°14'60.3 N, 152°01'29.1 E | 21 (17)<sup>m</sup> | 22 (16) |
| AN. Ant Atoll (East) | Pohnpei | 6°47'42.3 N, 158°01'20.7 E | 24 (16) | 22 (13) |
| RO. Roj | Pohnpei | 6°46'37.7 N, 158°12'24.1 E | 24 (16) | 23 (15) |
| CG. Coral Garden | Kosrae | 5°18'47.2 N, 162°53'01.8 E | 25 (15)<sup>m</sup> | 24 (13) |
| HP. Hiroshi Point | Kosrae | 5°15'88.0 N, 162°59'01.8 E | 25 (23)<sup>m</sup> | 25 (14) |
| **TOTAL** |  |  | **277 (190)** | **278 (178)** |

\* indicates that no individuals of this species were found
<sup>1</sup> indicates that individuals were not collected from this site but are likely present
<sup>m</sup> indicated sites where multiple *Symbiodinium* species were detected

**Table S4:**
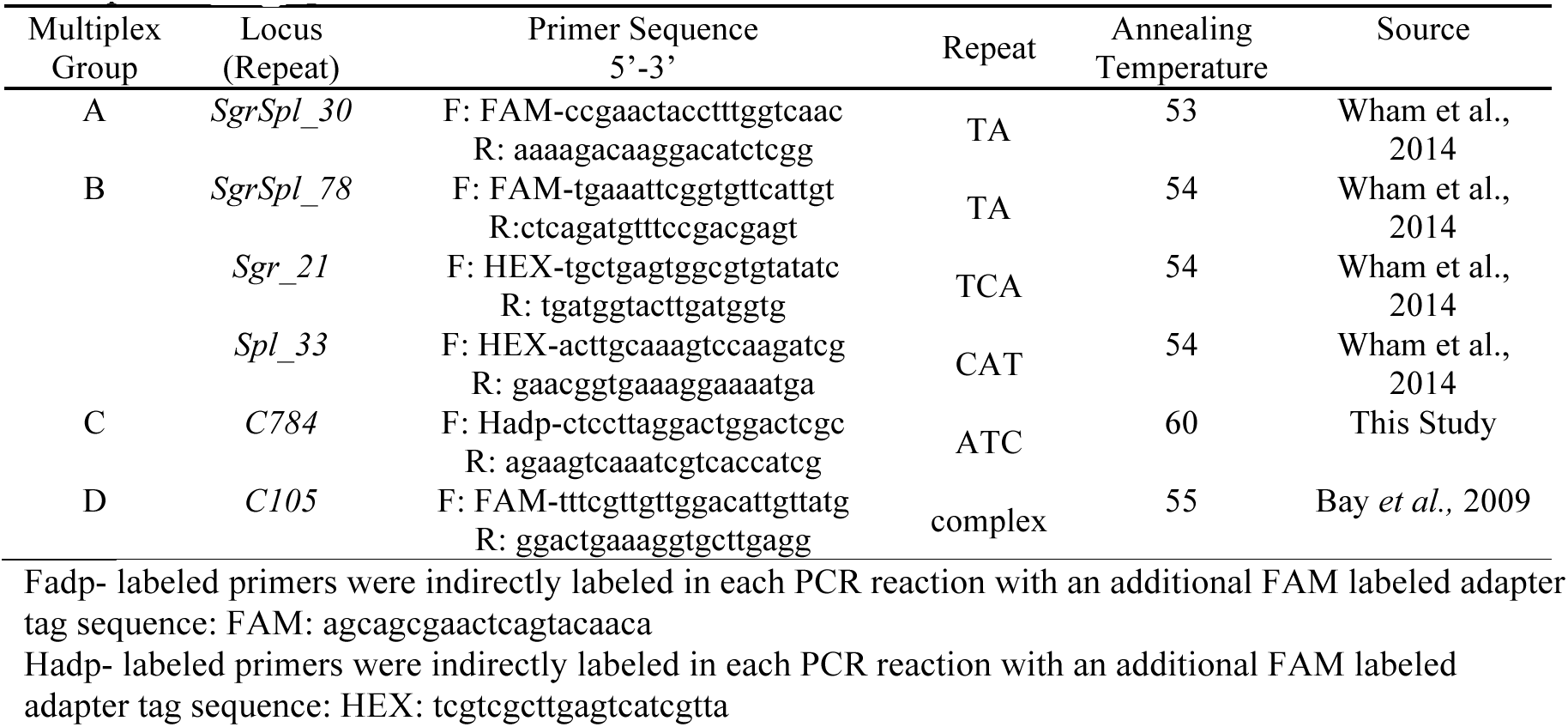
Summary of six polymorphic microsatellite loci used to assess genetic variation *Symbiodinium* clade C hosted by *A. hyacinthus* and *A. digitifera* and their corresponding multiplexing groups.

**Table S5:** Accession number and corresponding Clade C *Symbiodinium* lineage for each unique sequence identifier from Figure S2.

| Sequence ID | Accession Number | Lineage |
| --- | --- | --- |
| 1 | JQ043642 | C3 |
| 2 | KF572372 | C3 |
| 3 | JQ043643 | C3 |
| 4 | JQ043641 | C3 |
| 5 | JQ043640 | C3 |
| 6 | JQ043644 | C3 |
| 7 | JQ043635 | C3 |
| 8 | JQ043638 | C3 |
| 9 | JQ043637 | C3 |
| 10 | JQ043636 | C3 |
| 11 | KF572331 | C3 |
| 12 | KF572332 | C3 |
| 13 | KF572318 | C3 |
| 14 | KF572319 | C3 |
| 15 | KF572373 | C3 |
| 16 | KF572374 | C3 |
| 17 | KF572358 | C3 |
| 18 | KF572368 | C40 |
| 19 | KF572359 | C40 |
| 20 | KF572367 | C40 |
| 21 | KF572366 | C40 |
| 22 | KF572360 | C40 |
| 23 | KF572357 | C40 |
| 24 | KF572363 | C40 |
| 25 | KF572365 | C40 |
| 26 | KF572364 | C40 |
| 27 | KF572370 | C40 |
| 28 | JQ043603 | C31 |
| 29 | JQ043599 | C31 |
| 30 | JQ043604 | C31 |
| 31 | JQ043671 | C27 |
| 32 | JQ043673 | C27 |
| 33 | JQ043670 | C27 |
| 34 | JQ043669 | C27 |
| 35 | JQ043676 | C27 |

